# Scaling of optogenetically evoked signaling in a higher-order corticocortical pathway in the anesthetized mouse

**DOI:** 10.1101/154914

**Authors:** Xiaojian Li, Naoki Yamawaki, John M. Barrett, Konrad P. Körding, Gordon M. G. Shepherd

## Abstract

Quantitative analysis of corticocortical signaling is needed to understand and model information processing in cerebral networks. However, higher-order pathways, hodologically remote from sensory input, are not amenable to spatiotemporally precise activation by sensory stimuli. Here, we combined parametric channelrhodopsin-2 (ChR2) photostimulation with multi-unit electrophysiology to study corticocortical driving in a parietofrontal pathway from retrosplenial cortex (RSC) to posterior secondary motor cortex (M2) in mice *in vivo*. Ketamine anesthesia was used both to eliminate complex activity associated with the awake state and to enable stable recordings of responses over a wide range of stimulus parameters. Photostimulation of ChR2-expressing neurons in RSC, the upstream area, produced local activity that decayed quickly. This activity in turn drove downstream activity in M2 that arrived rapidly (5-10 ms latencies), and scaled in amplitude across a wide range of stimulus parameters as an approximately constant fraction (~0.2) of the upstream activity. A model-based analysis could explain the corticocortically driven activity with exponentially decaying kernels (~20 ms time constant) and small delay. Reverse (antidromic) driving was similarly robust. The results show that corticocortical signaling in this pathway drives downstream activity rapidly and scalably, in a mostly linear manner. These properties, identified in anesthetized mice and represented in a simple model, suggest a robust basis for supporting complex non-linear dynamic activity in corticocortical circuits in the awake state.

**SIGNIFICANCE STATEMENT:** The signaling properties of corticocortical connections are not well understood, particularly for higher-order inter-areal pathways. Here, we developed a paradigm based on parametric optogenetic photostimulation, linear-array electrophysiology, and mathematical modeling to characterize signaling along corticortical connections linking retrosplenial cortex to posterior secondary motor cortex (M2) in anesthetized mice. The results indicate that corticocortically driven activity in the downstream area followed the optogenetically evoked upstream activity in a rapid and scalable manner, and could be described with a simple linear integrator model. These findings suggest that this pathway, when activated selectively in the unconscious state, supports intrinsically linear inter-areal communication.

## INTRODUCTION

Corticocortical pathways support inter-areal communication, which is central to behavior (Felleman and Van Essen, 1991; Misic and Sporns, 2016). Quantitative characterization of signaling in corticocortical pathways is essential for understanding and modeling how they contribute to information-processing. This information can also help to address a fundamental question in connectomics research, of how the relatively static structure of corticocortical networks can give rise to the complex non-linear dynamic activity typically observed in awake animals (Park and Friston, 2013). For example, it is unknown whether such non-linearities are present already at the most basic level of the intrinsic biophysical properties of corticocortical connections, or whether they arise at higher levels of network interactions.

Connectomics studies have identified a structural basis for many corticocortical pathways (Oh et al., 2014; Zingg et al., 2014; Jbabdi et al., 2015), and optogenetic mapping studies have begun to characterize dynamic signaling at the mesoscopic scale (Lim et al., 2012). However, the properties of inter-areal signaling in these pathways have been challenging to resolve *in vivo*, particularly in higher-order pathways, which are many synapses removed from the sensory periphery and thus difficult to analyze in an isolated, selective, and spatiotemporally precise manner with natural stimuli. Extracellular electrical stimulation has been used in efforts to artificially generate focal activity, but is inherently limited due to its nonspecificity, antidromic activation, and other issues (Nowak and Bullier, 1998; Histed et al., 2009).

Recently developed optogenetic methods hold promise for overcoming these limitations. Such methods have enabled detailed characterization of cell-type-specific connections in long-range circuits *ex vivo* (Petreanu et al., 2007; Petreanu et al., 2009). Corticocortical circuits in mice have begun to be characterized at the cellular level with this approach (Mao et al., 2011; Hooks et al., 2013; Yang et al., 2013; Kinnischtzke et al., 2014; Petrof et al., 2015; Suter and Shepherd, 2015; Kinnischtzke et al., 2016; Sreenivasan et al., 2016), mostly focusing on lower-order corticocortical pathways that involve primary cortical areas. In parallel are efforts to use similar approaches *in vivo* to characterize how optogenetically evoked activity interacts with sensory input at the level of primary sensory cortex (e.g. (Manita et al., 2015; Reinhold et al., 2015)).

Among these newly characterized corticocortical circuits, however, is a higher-order projection from retrosplenial cortex (RSC) to posterior secondary motor cortex (M2) (Yamawaki et al., 2016). RSC axons innervate M2 neurons broadly across all layers and projection classes, forming a synaptic circuit whereby RSC, which receives input from dorsal hippocampal networks and is involved in spatial memory and navigation, appears to communicate with M2, which sends output to diverse motor-related areas and appears to be involved in diverse sensorimotor functions. As such, this connection is an interesting target for the reverse engineering of signaling properties in a higher-order inter-areal corticocortical pathway.

Here we have developed an approach based on combining the slice-based circuit analysis methods (Yamawaki et al., 2016) with system-identification methods used in sensory systems research (Wu et al., 2006) to develop an *in vivo* paradigm suitable for assessing and manipulating corticocortical circuit dynamics in the intact brain. We used the same labeling paradigms to express ChR2 in presynaptic RSC neurons, and developed *in vivo* methods in the ketamine-anesthetized mouse for sampling photo-evoked multi-unit activity in M2 driven by RSC photostimulation. Duplication of the setup to permit both stimulation and recording at both ends of the RSC→M2 projection allowed a detailed parametric characterization of both local (upstream) and downstream activity evoked both ortho- and antidromically. This allowed us to systematically investigate how optogenetically evoked RSC→M2 signaling drives downstream activity as a function of upstream stimulation amplitude and duration. The parametric nature of the data collected with this approach allowed us to also assess the linearity of corticocortical signaling in this pathway.

## MATERIALS AND METHODS

### Animals

Studies were approved by the Northwestern University Animal Care and Use Committee, and followed the animal welfare guidelines of the Society for Neuroscience and National Institutes of Health. Wild-type mice (*Mus musculus*, C57BL/6, female and male; Jackson Laboratory, Bar Harbor, ME) were bred in-house. Mice were 6-9 weeks old at the time of *in vivo* experiments.

### Stereotaxic injections

Mice under deep anesthesia underwent stereotaxic injection of adeno-associated virus (AAV) carrying ChR2 into the RSC, following standard methods as previously described (Yamawaki and Shepherd, 2015; Yamawaki et al., 2016). Viruses used were: AAV1.CAG.ChR2-Venus.WPRE.SV40 (AV-1-20071P, University of Pennsylvania Vector Core, Philadelphia, PA; Addgene #20071, Addgene, Cambridge, MA), and AAV9.CamKIIa.hChR2(H134R)-eYFP.WPRE.hGH (AV-9-26969P, Penn Vector Core; Addgene #26969P). Stereotaxic coordinates for the RSC were: −1.4 mm caudal to bregma, ~0.5 mm lateral to midline. To minimize cortical damage, the glass injection pipette was pulled to a fine tip, beveled to a sharp edge (Micro Grinder EG-400, Narishige, Tokyo, Japan), and advanced slowly into the cortex; injections were made slowly (over 3 minutes) at two depths (0.8 and 1.2 mm from pia, ~20 nL per injection). Mice were returned to their home cages and maintained for at least 3 weeks prior to experiments, to allow time for ChR2 expression levels to rise in the infected neurons.

### Cranial hardware

Mice under deep anesthesia underwent placement of cranial mounting hardware. A small skin incision was made over the cerebellum to expose the skull, and a stainless-steel set screw (single-ended #8-32, SS8S050, Thorlabs, Newton, NJ), crimped with a spade terminal (non-insulated, 69145K438, McMaster-Carr, Elmhurst, IL), was affixed with dental cement (Jet Denture Repair Powder, Lang Dental Manufacturing Co., Inc., Wheeling, IL) to the skull. This set screw was later screwed into the tapped hole located at the top of a 1/2″ optical post (Thorlabs) used for head fixation.

### *In vivo* circuit analysis: general procedures

Mice were anesthetized with ketamine-xylazine (ketamine 80-100 mg/kg and xylazine 5-15 mg/kg, injected intraperitoneally), placed in the recording apparatus, and head-fixed using the set screw as described above. Body temperature was monitored with a rectal probe and maintained at ~37.0 °C via feedback-controlled heating pad (FHC, Bowdoin, ME). Craniotomies were opened over the RSC and M2 using a dental drill (EXL-M40, Osada, Los Angeles, CA), just large enough (~1 mm) to allow passage of the linear arrays and the tips of the optical fibers. During the subsequent recordings, ACSF (containing, in mM, 127 NaCl, 25 NaHCO_3_, 1.25 NaH_2_PO_4_, 2.5 KCl, 25 D-glucose; all reagents from Sigma-Aldrich, St Louis, MO) was frequently applied to the exposed brain area to prevent damage from dehydration. The level of anesthesia was continuously monitored based on paw pinching, whisker movement, and eye-blinking reflex. Additional doses of anesthesia were given subcutaneously (50% of induction dose) when required.

### Photostimulation apparatus

An optical fiber (FG400AEA, multimode fiber, 0.22 NA, 400 μm core, outer diameter 550 μm with coating; Thorlabs), mounted on a motorized micromanipulator (Sutter Instrument, Novato, CA), was positioned directly over the region of the infected neurons in the RSC. The tip of the fiber was ~0.5 mm away from the surface of the brain, immersed in ACSF. In most experiments, a second fiber was similarly positioned directly over the M2. For each fiber, the light source was an LED (M470L3, Thorlabs), coupled to the fiber by an adapter (SM1SMA, Thorlabs). The power was controlled using an LED driver (LEDD1B, Thorlabs; or, driver based on RCD-24-1.00 module, RECOM Lighting, Neu-Isenburg, Germany). The output power of the LED driver was modulated by signal waveforms delivered via a multifunction analog and digital interface board (NI USB 6229; National Instruments, Austin, TX) or by a signal generator based on a 32-bit microcontroller board (Arduino Due with ARM Cortex-M3, Adafruit, New York, NY). The boards were also used to send a short pulse train to digitally encode the start and other parameters of the light waveform, sampled on the digital input port of the electrophysiology data acquisition (DAQ) board. Software tools (LabVIEW) included a GUI (GenWave) for generating and transferring the waveforms to the LED controller. The LED driver was either internally software-triggered (GenWave) or externally hardware-triggered by a digital signal. A power meter was used to calibrate the relationship between input voltage to the driver and the output intensity of the fiber, and the calibration curve was used to determine the voltages (in the range of 0–5 V) corresponding to 0, 20, 40, 60, 80, and 100% of the full power (6.1 mW, measured at the tip of the optical fiber). During the experiment, analog voltages corresponding to these intensities were sent to the LED driver.

### Electrophysiology apparatus

The linear arrays used were 32-channel silicon probes with ~1 MΩ impedance and 50-μm spacing (model A1×32-6mm-50-177, NeuroNexus, Ann Arbor, MI), in either “triangular” or “edge” configuration. The probes were mounted on a motorized 4-axis micromanipulator (Thorlabs MTSA1 linear translator mounted on a Sutter MP285 3-axis manipulator), and positioned under stereoscopic visualization over the M2 at cortical surface (i.e., entry point) coordinates of +0.6 mm rostral to bregma and 0.2 mm lateral to midline. The probes were tilted by ~30° off the vertical axis for alignment with the radial axis of the cortex. The probe was then slowly inserted into the cortex at a rate of 2 μm/s (controlled by software), until it reached a depth of 1600 μm from the pia. In most experiments, a second array was similarly inserted into the RSC (same stereotaxic coordinates as given above for the viral injections), except that in this case the array was inserted perpendicular to the horizontal plane, and the optical fiber was slightly tilted.

Signals were amplified using an amplifier board based on a RHD2132 digital electrophysiology interface chip (Intan Technologies, Los Angeles, CA). The RHD2132 chip is an analog front end that integrates the analog instrument amplifiers, filters, analog-to-digital (ADC) converters, and microcontrollers in one chip. The serial peripheral interface (SPI) port is used to configure the chip and to stream the silicon probe data to the data acquisition (DAQ) board. The gain of the amplifier was fixed at 96 × 2 = 192 (2-stage amplifier). The filter was set to an analog bandpass of 0.1~ 7.5K Hz with a digital filter cutoff of 1 Hz. Because the 32 channels of the silicon probe inputs share the same 16 bit ADC with a multiplexer, and the maximum sample rate of the ADC is 1.05 x 10^6^ samples per second (SPS), the single channel sample rate was set to 30,000 SPS.

For hardware control, we used a RHD2000 USB Interface Evaluation Board (Intan) or DAQ board based on a breakout board with a XEM6010 USB/FPGA module (Opal Kelly, Portland, OR), a field-programmable gate array (FPGA) with many digital I/O channels for communicating with other digital devices and streaming in the linear array data from the RHD2000 amplifiers. The USB port of the module was linked with a USB cable to pipe the data stream in or out of the PC. The RHD2000 amplifier boards were connected to a DAQ board using SPI interface cables in low-voltage differential signal mode, which is well suited for communication via longer cables. In this experiment, the digital ports included in the DAQ board were only used for acquiring the LED photostimulation parameters from the LED controller.

For data logging, The C++/Qt based experimental interface evaluation software (Intan) was used for early stage evaluation. Then the original APIs (Rhythm USB/FPGA interface) were all rebuilt and wrapped up into a LabVIEW-based SDK. All the software, including the amplifier configuration, online visualization, data logging, and more, were developed from this SDK in LabVIEW environment.

### Trace acquisition and analysis

With the mouse anesthetized and head-fixed and the two linear arrays and two optical fibers in place, photostimuli were repeatedly delivered while continuously sampling electrophysiological activity across the 32 channels per array. In each trial, a single photostimulus was delivered on one fiber, using one of the 25 combinations of stimulus intensities (20, 40, 60, 80, or 100 percent of maximum) and durations (1, 5, 10, 20, or 50 ms). Across all trials, all 25 stimulus combinations were tested, in randomly interleaved sequence, with an inter-trial (interstimulus) interval of 2 sec, This cycle was repeated many (e.g. ~30) times, and the entire process was repeated again for the second optical fiber (if present). For stimulation on each fiber, the resulting electrophysiological data set typically consisted of 2 arrays x 32 channels/array x 25 stimulus parameter combinations x 30 trial repetitions = 48,000 traces.

These trace data were stored as the raw signal from the amplifiers, and filtered as follows. A digital 60 Hz notch filter (Matlab) was used to reduce line hum. A digital high-pass filter (800 Hz cut-off, 2^nd^-order Butterworth; Matlab) was used to isolate the higher-frequency components of the electrophysiology responses for event detection.

For event detection, in this study we focused on analyzing multi-unit activity; although single units could be isolated on some channels, single-unit analysis was generally hampered by the short-latency barrage of activity just after a photostimulus, particularly in the upstream area and especially at higher stimulus intensities. Similar to previous studies of multi-unit activity, we defined “events” (i.e., spikes in the traces) as voltage excursions that were ≥4 standard deviations (s.d.) above the baseline amplitude (measured in the 1500 ms prior to the stimulus), lasting ≥0.1 ms (3 continuous samples) in duration. Event detection based on these criteria was performed on all traces using Matlab routines.

Peristimulus time histograms were constructed as follows, using Matlab (Mathworks, Natick, MA) routines. For each trial, time stamps were determined for each detected event, and the time stamps of all the events of every channel were used to generate a single-trial raster plot using 1-ms bins (**Fig. 1G, top**). Trials were repeated multiple times, and raster plots were grouped by experimental condition (e.g. each particular stimulus parameter combination) and averaged over all trials (typically ~30 trials) to obtain a multi-trial histogram showing the mean activity across all channels for that condition (**Fig. 1G, middle**). The multi-trial histograms were also summed across channels to obtain an all-channel histogram for each condition (**Fig. 1G, bottom**).

**Figure 1.**
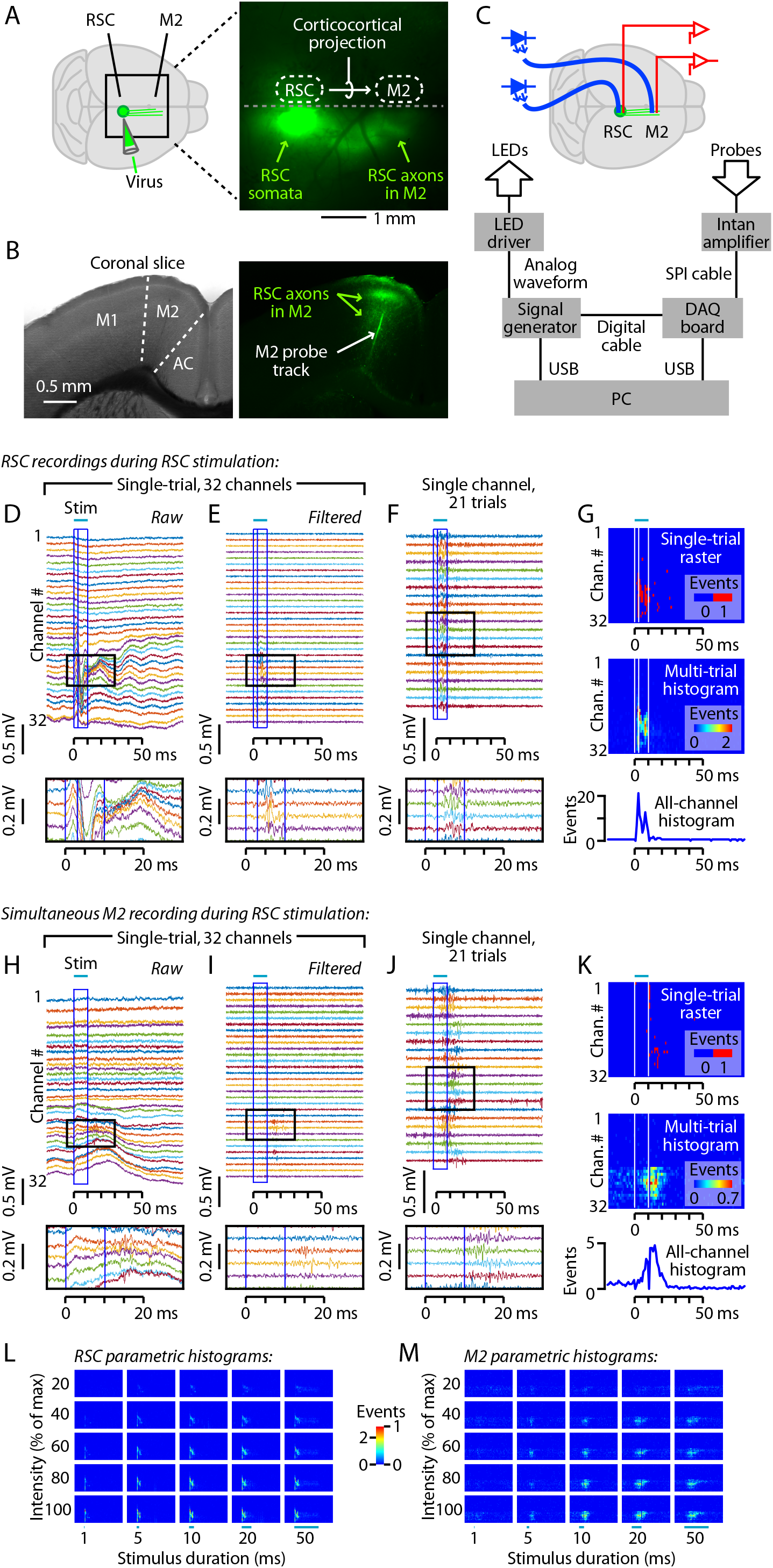
Experimental paradigm for characterizing inter-areal signaling in the corticocortical projection from retrosplenial (RSC) to posterior secondary motor (M2) cortex. (A) Virus injection in RSC infects somata at the injection site, resulting in anterograde labeling of RSC axons projecting to M2. Right: epifluorescence image of the dorsal surface of the brain of an anesthetized mouse, showing labeled axons projecting from RSC to posterior M2. (B) Coronal brain slices showing labeled axons in M2, and the track of a dye-coated linear array. Left: bright-field image. M2 is between the primary motor (M1) and anterior cingulate (AC) cortices. Right: epifluorescence image, showing labeled axons from RSC within M2, and the track of a dye-coated linear array (probe) that had been inserted in M2. (C) Depiction of experimental set-up showing aspects of the hardware control apparatus and wiring. An optical fiber (blue) was placed over, and a silicon probe was inserted into, each of the two cortical areas. The optical fibers were coupled to blue light-emitting diodes (LEDs). For clarity, only the fiber over the RSC is depicted here. See Methods for additional details. (D-G) Examples of RSC recordings during RSC photostimulation. (D) Raw (unfiltered) traces from the 32-channel linear array in the RSC, recorded during a single trial of RSC photostimulation (10 ms pulse, 100% intensity). The stimulus is indicated by the bar above, and by the blue box. The interior line with in the box indicates the 3-ms post-stimulus time point, which was the maximal width of the responses that were blanked to eliminate a stimulus artifact. The region demarcated by the black box is shown in the inset at the bottom. (E) Same, but after high-pass filtering. (F) Traces from a single channel, recorded on multiple stimulus presentations. Photostimulation reliably generated post-stimulus activity. (G) Top: Raster plot of detected events for a single trial (traces shown in E). Middle: Histogram showing events detected across all trials for each channel. Bottom: Overall histogram, calculated by summing across all channels. (H-K) Same as D-G, for the recordings made simultaneously from the linear array inserted in M2. (L) Histograms for the RSC recordings, for 25 combinations of stimulus durations (1, 5, 10, 20, and 50 ms) and intensities (20, 40, 60, 80, and 100% of maximum). (M) Same, for the M2 recordings.

The raw traces were contaminated by a brief stimulus artifact immediately after stimulus onset and offset. These transients were greatly attenuated by digital high-pass filtering (described above). The duration of the residual transients was estimated for each experiment (i.e., animal), and ranged from 1 to 3 ms. For display, both transients were simply blanked for this brief duration. For subsequent analyses, the data were replaced in the following way, taking advantage of the availability of responses recorded using different stimulus durations. For the onset transient, the event count of the blanked window was replaced by the average value of the baseline window over the 20 ms pre-stimulus interval. For the offset transient, the event count of the blanked window was replaced by the event count measured for the next-longer-duration stimulus acquisition. For example, in the case a 2-ms-long transient in the responses recorded during a 10-ms-long stimulus, the data over the interval of 11-12 ms post-stimulus (i.e., the 11^th^ and 12^th^ 1-ms bins) were replaced by the data value recorded at 11-12 ms during the 20-ms-long stimulus, and so on. However, for the longest-duration stimulus of 50 ms, we instead replaced with the baseline values, as post-stimulus activity had returned to approximately baseline levels by this time.

### Laminar analysis

We estimated the depth of probe insertion in the cortex (and thus the cortical depth of each contact) based on the total displacement of the motorized manipulator holding the probe. In addition, because this estimate can be affected by the viscoelastic properties of brain tissue, we also routinely analyzed the electrophysiological traces to estimate the depth of insertion. For this, we calculated variance in the FFT of the voltage traces to identify the transition from low-variance exterior channels and high-variance intracortical channels. The estimated depth based on this approach matched well with the estimated depth based on images of the electrode at the site of penetration into the brain. Using this combination of approaches, the estimated probe depths were thus likely to be accurate within 50-100 μm. Additionally, in a subset of experiments, probe tracks were labeled by coating the probe with fluorescent dye, and subsequently visualized in brain slices with epifluorescence optics to verify accurate placement of the probes in the M2 and/or RSC.

### Model based analysis

We fit the following model to the locally evoked activity in RSC:

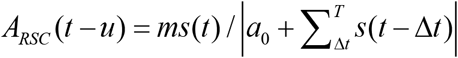

where *m* is a scaling factor, *a*_0_ regulates the strength of decay, Δ*t* indexes the delays over which stimulation affects activity (Δ*t* = 0 would be instantaneous activation), *s*(*t*) is the optical stimulus and *u* is the delay. The three parameters of this model *u*, *m*, and *a*_0_ are optimized to minimize the root mean squared error (RMSE) of the model using the MATLAB fminsearch function.

We fit the following model to the downstream activity in M2:

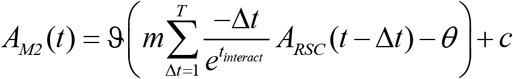

where *m* is a scaling factor, 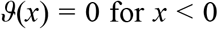 for *x* < 0 and 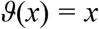 for *x* > 0, Δ*t* indexes the past input from RSC, *τ_interact_* is the interaction time constant, *A_RSC_*(*t*) is the activity in area RSC, *θ* is the threshold, and *c* is the baseline. The four parameters of this model *c*, *θ*, *τ_interact_*, and *m* are also optimized to minimize the RMSE.

### Experimental Design and Statistical Analysis

The main data set comes from an experimental design involving parametric stimulation and recording that will be explained in detail in the Results. In general, unless otherwise stated, the following statistical methods were used. Descriptive statistics are reported and displayed as sample medians ± median absolute deviations (m.a.d.) (calculated with the Matlab function, mad.m). Group data are compared using appropriate non-parametric tests (e.g. rank sum tests for unpaired and sign tests for paired data) as indicated, with significance defined as *p* < 0.05.

## RESULTS

### RSC photostimulation drives downstream M2 activity

To investigate corticocortical signaling in the RSC→M2 pathway, we used viral methods to label the RSC neurons with ChR2, optical fibers to photostimulate them, and linear arrays to record the evoked activity. Similar to previous studies of this pathway (Yamawaki et al., 2016), we infected neurons in RSC with an AAV encoding ChR2 and a fluorescent protein (**Fig. 1A,B**). After a recovery period of several weeks, animals were anesthetized with ketamine and underwent placement of a photostimulation fiber over the RSC and silicon probes in both the RSC and M2 (**Fig. 1C**). (As described at the end of the Results, a second optical fiber was also routinely placed over the M2 to enable antidromic activation; however, the main focus of the study is on the ‘forward’ orthodromic signaling evoked by RSC stimulation.)

With this optogenetic photostimulation and electrophysiological recording arrangement, we photostimulated ChR2-expressing neurons in RSC and sampled responses simultaneously in RSC (**Fig. 1D-G**) and M2 (**Fig. 1H-K**). The raw traces (**Fig. 1D,H**) were high-pass filtered (**Fig. 1E,I**), revealing brief barrages of photo-evoked events on multiple channels on both probes, easily discernable in single trials. Over repeated trials, photostimulation reliably evoked spiking activity on channels showing responses (**Fig. 1F,J**). We analyzed the traces to detect events, representing multi-unit activity (see Methods), and used the timing of events to construct single-trial rasters and multi-trial peristimulus time histograms (**Fig. 1G,K**). These histograms showed robust, transient increases in multi-unit activity starting with a short delay after the onset of photostimulation in RSC, for both the RSC and M2 recordings. The example traces and histograms are for responses to a photostimulus with 10 ms duration and maximal intensity, extracted from a much larger data set using 25 different combinations of stimulus durations and intensities (**Fig. 1L,M**).

Prior to presenting our analyses of these parametric data sets in detail in later sections, we present some additional characterizations of the technique. One consideration is whether responses differ for different viruses and constructs, and we therefore performed parallel experiments with two different AAV serotypes carrying different variants of ChR2 driven by different promoters: AAV1-ChR2-Venus, carrying wild-type ChR2 driven by the CAG promoter, and AAV9-ChR2-eYFP, carrying ChR2 with the H134R mutation driven by the CaMKII promoter (see Methods). The two viruses gave very similar responses (as described in later sections), suggesting that our strategy is not overly affected by the particular types of viruses and opsin constructs used. Both viruses infected cortical neurons only locally at the injection site in the RSC, without evidence of retrograde infection in M2, as shown previously (Yamawaki et al., 2016); i.e., the M2 contained anterogradely labeled axons of RSC neurons, but not retrogradely labeled somata of M2 neurons.

The brief burst of multi-unit activity observed in M2 (**Fig. 1I-K**) arriving shortly after that in RSC (**Fig. 1E-G**) suggests that spiking activity in RSC neurons propagated via their corticocortical axons and synaptically drove spiking activity in M2 neurons, via the abundant excitatory RSC→M2 connections previously described for this corticocortical circuit (Yamawaki et al., 2016). Alternatively, events detected in M2 might represent spikes in presynaptic axons rather than in postsynaptic neurons. However, this seems unlikely, particularly as spikes in thin corticocortical axons are much smaller in amplitude and usually difficult to detect (Raastad and Shepherd, 2003). To assess whether the M2 responses primarily reflect synaptically driven spikes of postsynaptic M2 neurons, rather than spikes in presynaptic axons, we sampled M2 responses before and after injecting M2 with muscimol (100 nL, 5 mM in ACSF), a GABA agonist, which suppresses spiking in cortical neurons while preserving presynaptic spiking (Chapman et al., 1991; Chatterjee and Callaway, 2003). We also simultaneously recorded the activity in RSC, to control for the possibility that muscimol injected into M2 might diffuse to RSC. As expected, muscimol injection in M2 had no effect on activity in RSC but abolished most of the activity in M2 (**Fig. 2A;** 4 of 4 animals). A similar effect was observed when blockers of glutamatergic synaptic transmission (100 nL of 1 mM CNQX and 5 mM CPP in ACSF) were injected in M2 (1 animal). Pooling the muscimol and CNQX/CPP data showed no effect of the drugs on RSC activity but a significant effect on M2 responses (**Fig. 2B**; *p* = 0.013, *t*-test, *n* = 5). Injection of saline had no effect (2 of 2 animals). Thus, M2 responses appear to be primarily driven by corticocortical synaptic activity.

**Figure 2.**
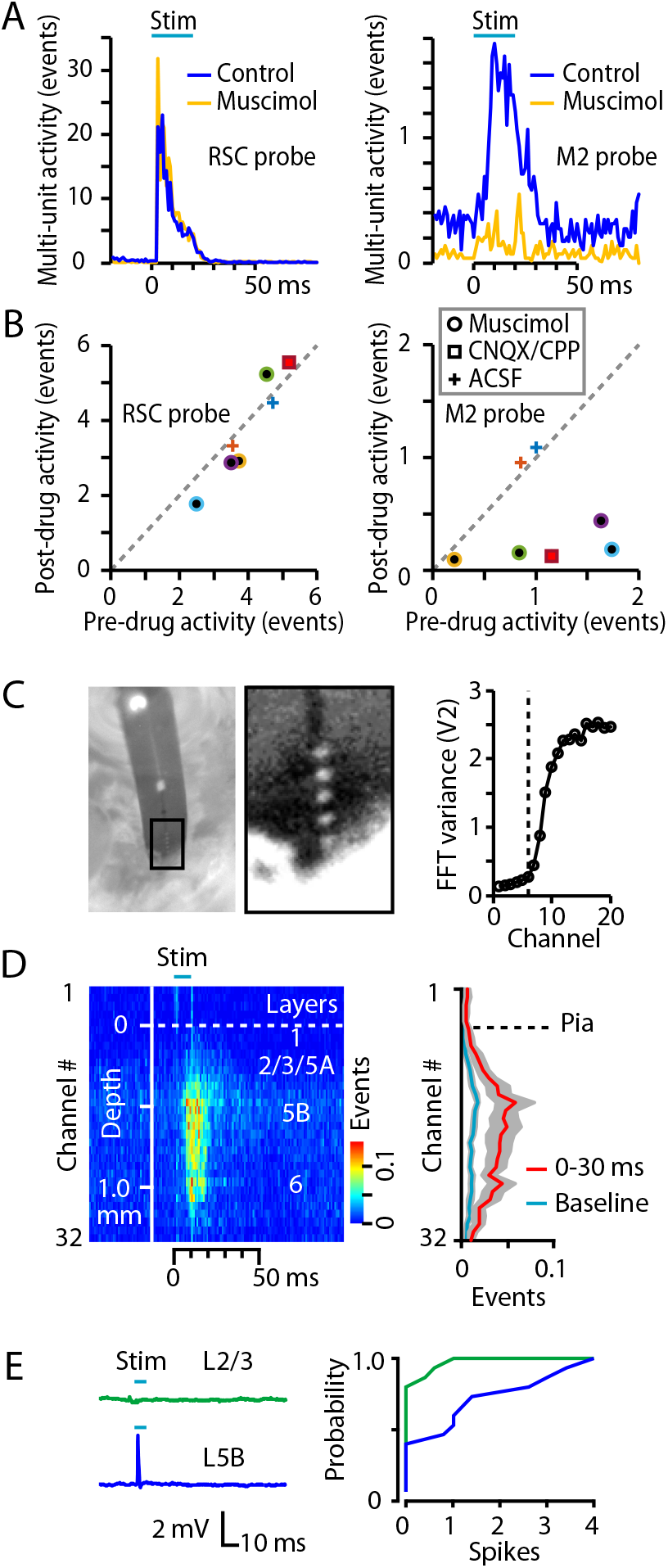
Additional characterizations of the technique. (A) Example histograms for responses to RSC stimulation, recorded before and after injection into M2 of muscimol. Muscimol, which suppresses synaptically driven activity, greatly reduced downstream activity in M2 (right) but not in RSC (left). (B) Responses recorded before and after injection into M2 of muscimol, CNQX/CPP, or ACSF (see legend for symbols). Drug injection did not affect activity in RSC (left) but significantly reduced responses in M2 (right; *p* = 0.013, *t*-test, *n* = 5). Colors indicate different experiments (animals). (C) Left: Image of 32-channel silicon probe, taken through the ocular of a stereoscope, showing 5 visible contacts above the penetration site into the cortex. Distance between contacts is 50 μm. Right: Plot of the variance in the FFT of the traces collected on the first 20 channels of the probe, showing an abrupt increase for channels deeper than the 6^th^ contact (dashed line). (D) Left: Average peristimulus-time histogram across all channels in a 32-channel array in M2 during RSC photostimulation, plotted on a color scale (mean ± s.e.m., for *n* = 9 mice injected with AAV1-ChR2). Right: Average laminar profile, plotted as the average event rate per channel during the response interval (red) and baseline (blue). (E) In *ex vivo* brain slice experiments, cell-attached recordings were made from layer 2/3 and layer 5B neurons while photostimulating RSC axons. Left: Example traces showing spiking response in the layer 5B neuron. Right: The mean number of evoked spikes was calculated for each neuron, and plotted as a cumulative histogram of spike probability. Layer 5B neurons spiked significantly more than layer 2/3 neurons (*p* = 0.009, rank-sum test; median spikes were 0 vs 1 for layer 2/3 vs 5B, respectively; *n* = 15 layer 2/3 and 15 layer 5B neurons).

We considered the sensitivity of the results to probe placement. In earlier pilot experiments the probe was sometimes inadvertently placed slightly lateral by ~0.5-1 mm, resulting in recordings in M1 instead of M2. In this case we observed little or no photo-evoked activity, consistent with the anatomy and electroanatomy of the RSC→M2 projection, which provides little or no direct input to M1 neurons (Yamawaki et al., 2016). Thus, accurate probe placement is important, but inaccurate placement would simply decrease the observed activity.

We also considered the laminar profile of M2 activity. To estimate the depth of penetration of the silicon probes (32 channels and 50 μm spacing), they were inserted leaving ~5 contacts out of the cortex, as verified both by viewing the site of entry with a high-power stereoscope, and assessing channel noise variance, which was low for contacts outside cortex (see Methods) (**Fig. 2C**). Group analysis (n = 9 mice injected with AAV1-ChR2) indicated wide distribution of activity across channels, and thus cortical layers, albeit with a bias towards middle and deeper layers (**Fig. 2D**). Previous slice-based characterization of RSC→M2 connectivity indicated that RSC axons form monosynaptic excitatory synapses onto postsynaptic M2 neurons across all layers and major classes of projection neurons, including upper-layer neurons (Yamawaki et al., 2016). Because those experiments were performed in whole-cell voltage-clamp mode, here, to explore the cellular basis for the relatively lower activation of upper layers in M2 we performed similar brain slice experiments but with cell-attached current-clamp recordings, allowing assessment of the efficacy of RSC inputs in generating suprathreshold (spiking) activity in M2 neurons. Comparison of layer 2/3 and layer 5 neurons showed significantly greater tendency of photo-activated RSC axons to generate spikes in layer 5 neurons (**Fig. 2E**), consistent with the laminar profile recorded *in vivo*.

From the results of these initial characterizations we conclude that (i) optogenetically stimulating RSC drives a delayed, brief wave of spiking activity in M2; (ii) the evoked activity appears to reflect mostly the properties of the corticocortical circuit itself rather than the those of the viruses and/or constructs; (iii) the M2 activity appears to arise from orthodromically driven signaling along the RSC→M2 corticocortical pathway, rather than non-specific (e.g. cortex-wide) activation; and (iv) RSC drives M2 neurons across multiple layers, particularly the middle and deeper layers. Next, we turned to a more in-depth analysis of recordings made simultaneously in both cortical areas.

### Comparison of local RSC and downstream M2 activity evoked by RSC photostimulation

Recording simultaneously from both the RSC and M2 during RSC photostimulation allowed us to assess both the locally driven activity in upstream RSC and the orthodromically driven activity in downstream M2 (**Fig. 3A**). For clarity, here we present only the data obtained using a stimulus of 100% intensity and 10 ms duration. With RSC photostimulation the activity recorded locally in RSC rose rapidly at the onset of photostimulation, peaking at approximately 5 ms, and declined rapidly as well (**Fig. 2B**). That the peak response occurred shortly after the brief post-stimulus blanking interval (from 0 to maximally 3 ms, depending on the experiment; used to eliminate a photovoltaic transient, as described in Methods) suggests that the blanking procedure affected primarily the rising phase of the response waveform. Activity recorded simultaneously in M2 (**Fig. 2C**) followed with a brief latency (7.5 ms after the RSC peak for AAV9, and 6.5 ms for AAV1; **Fig. 2D,E**) and rose to lower levels than observed in RSC (RSC/M2 amplitude ratio: 3.8 for AAV9, 4.1 for AAV1; **Fig. 2F**).

**Figure 3.**
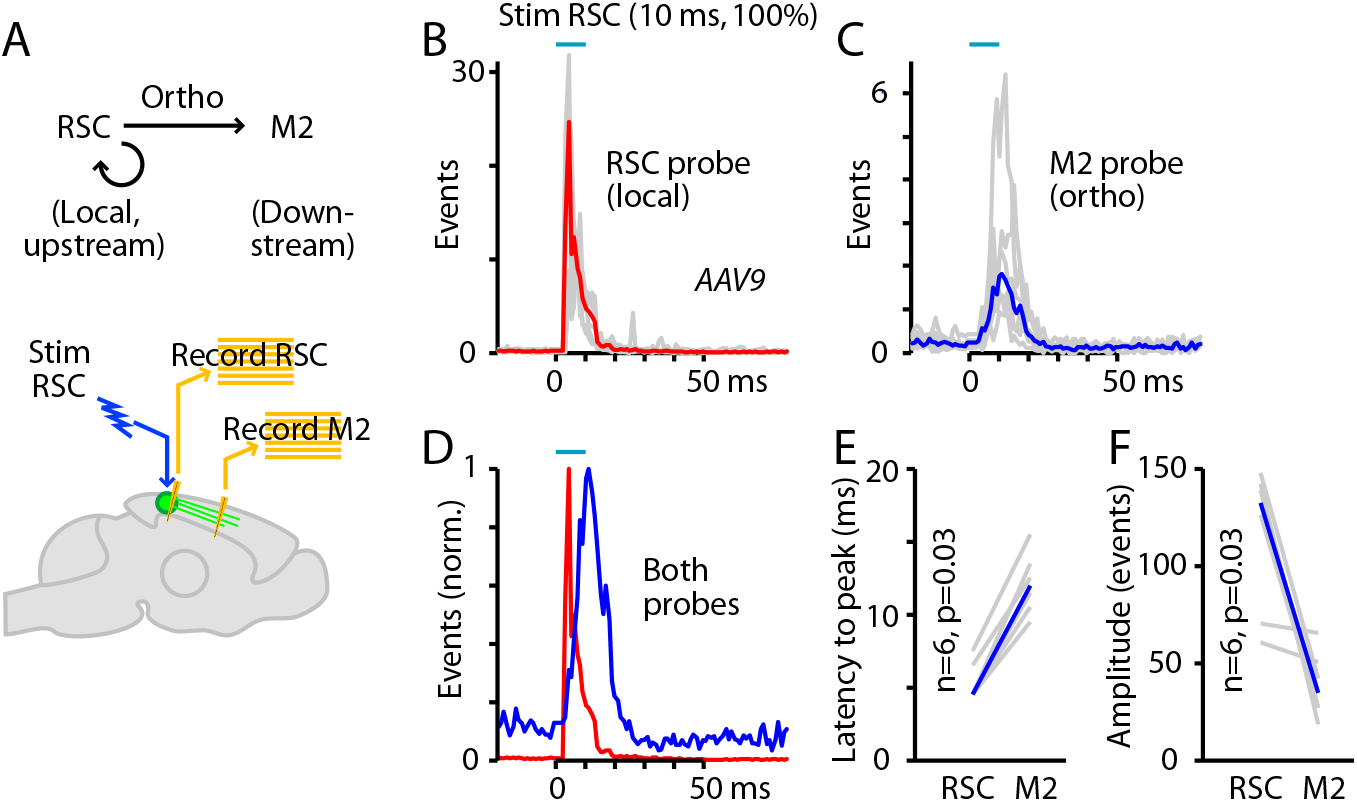
Comparison of local RSC and downstream M2 activity evoked by RSC photostimulation. (A) Experimental paradigm: RSC neurons were infected with AAV to express ChR2, and photostimuli were applied to RSC while recording multi-unit activity in both M2 (orthodromically driven) and RSC (locally driven). (B) Activity recorded on the RSC probe during RSC stimulation in animals injected with AAV9-ChR2. Red trace is the median response across 6 animals (traces for each animal shown in gray). (C) Activity recorded on the M2 probe during the same experiment. Blue trace is the median response across animals. (D) Overall activity on the RSC and M2 probes plotted together (peak-normalized). (E) Latencies (to peak) for responses recorded on the RSC and M2 probes. P-value calculated by 2-sided, paired sign test. (F) Amplitudes of responses (summed events) recorded on the RSC and M2 probes, plotted for each experiment (gray) and as the median across animals (blue). P-value calculated by 2-sided, paired sign test.

The results of this two-probe characterization of RSC photostimulation thus reveal two important aspects of corticocortical driving in this pathway. First, at the upstream end there is a rapid and strong decay of the local activity in the directly photostimulated RSC (**Fig. 2B,G**). The time course and extent of this decay are consistent with ChR2 desensitization (Nagel et al., 2003; Nagel et al., 2005; Lin et al., 2009), although other factors such as activation of interneurons and activity-dependent synaptic depression are also likely to contribute. Second, at the downstream end, the corticocortically driven activity in M2 was reduced in amplitude and slightly delayed relative to the RSC activity. A caveat is that these properties might not be generalizable, reflecting instead the particular photostimulus parameters used in these experiments. Therefore, we next investigated in detail the stimulus dependence of the responses by exploring a wide range of stimulus intensities and durations.

### Parametric characterization of orthodromic (forward) driving

Key parameters for the dynamics of a circuit are the dependency on stimulus amplitude (light intensity) and duration (pulse width). Stimulus trials were delivered at five different intensities (20, 40, 60, 80, and 100% relative to maximum) and durations (1, 5, 10, 20, and 50 ms), with random interleaving and many repetitions (typically ~30 trials per experiment) for each of the 25 unique intensity-duration combinations (Fig. 4A). Responses were averaged across trials as before, and the median responses on the local RSC probe (**Fig. 4B**) and the downstream M2 probe (**Fig. 4C**) were determined across animals. Clearly, the evoked activity in both RSC and M2 varied with stimulus parameters. To assess how response properties might depend systematically on stimulus parameters, we developed a simple model, and performed several further analyses.

**Figure 4.**
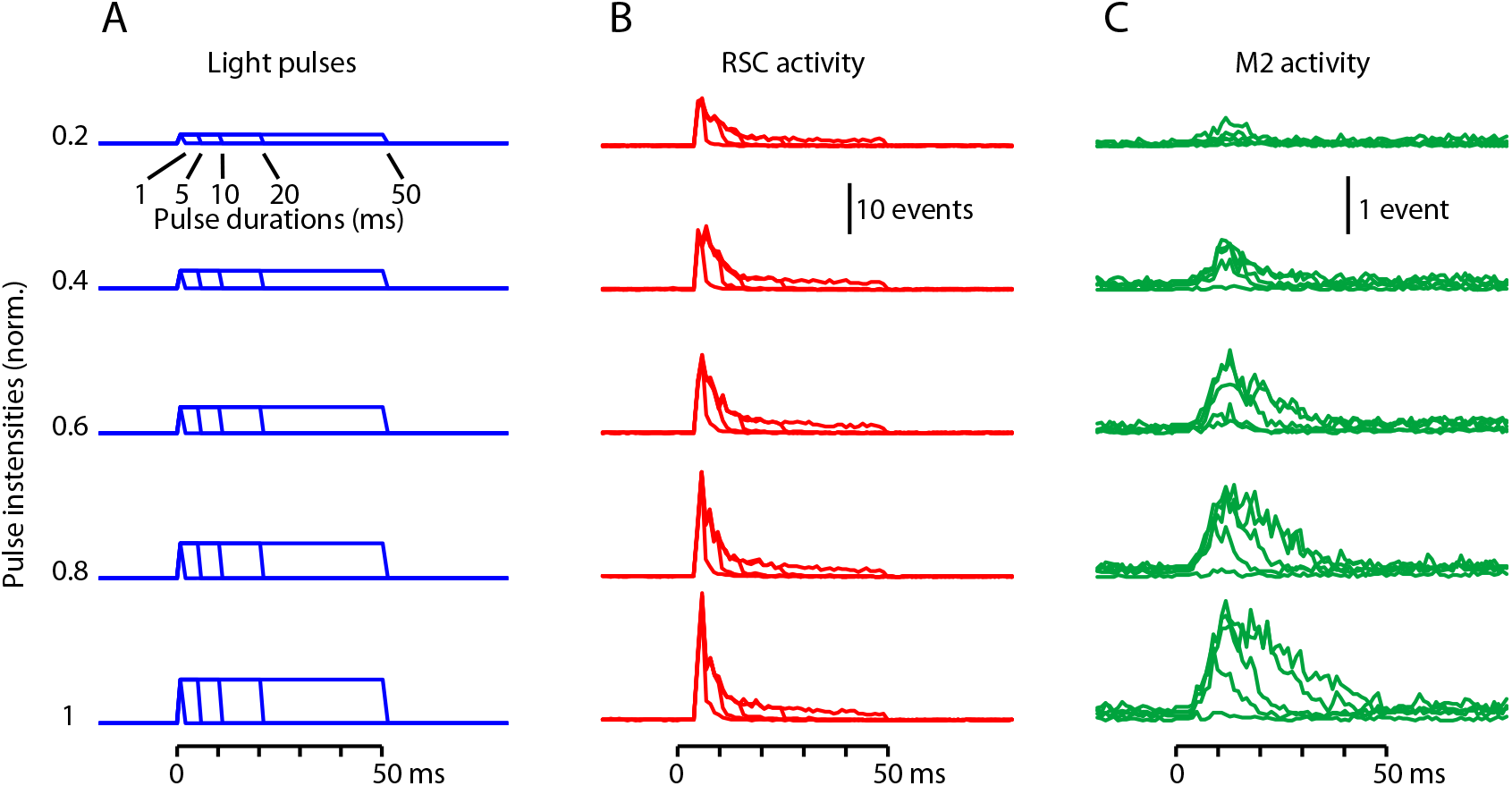
Parametric characterization of orthodromic (forward) driving. (A) Light pulses with a total of 25 different combinations of stimulus intensities (20, 40, 60, 80, and 100% relative to maximum) and durations (1, 5, 10, 20, and 50 ms) were used to photostimulate the RSC. (B) Activity recorded locally in RSC (red) in response to RSC photostimulation using the stimuli shown in panel A. Each trace is the median response across AAV9-ChR2 animals (*n* = 6 experiments). (C) Activity recorded simultaneously in M2 (green) in the same experiments.

### A simple two-stage model captures the major features of orthodromic driving

Visual inspection of the waveforms of both the RSC and M2 responses (**Fig. 4B,C**) showed roughly linear increases with intensity. Clearly, activity in the photostimulated RSC decays rapidly, consistent with ChR2 densensitization (as discussed above). However, in the downstream M2, it is unclear how responses scale with upstream RSC activity; for example, do they scale linearly, or show signs of adaptation? We would like a simple model to allow us to both describe and interpret the data.

Explorative data analysis revealed that we could fit the directly stimulated (upstream) area well with briefly delayed activation followed by a large and rapid decay (**Fig. 5A**). Hence, we modeled the response as a time-shifted delta function divided by a linear function of the integral of the stimulus history. So this first-stage model has 3 parameters for gain, delay, and the steady-state adaptation. These parameters seem intuitively necessary: the gain describes the strength of the locally generated activity; the delay is needed due to the ~3 ms blanking of the stimulus artifact (see Methods), but can also account for the kinetics of ChR2 activation and spike generation; and some degree of adaptation (of the locally generated activity) is expected from ChR2 inactivation/desensitization kinetics, and allows for additional factors contributing to a temporal decline in activity (e.g. GABA release, synaptic depression).

**Figure 5.**
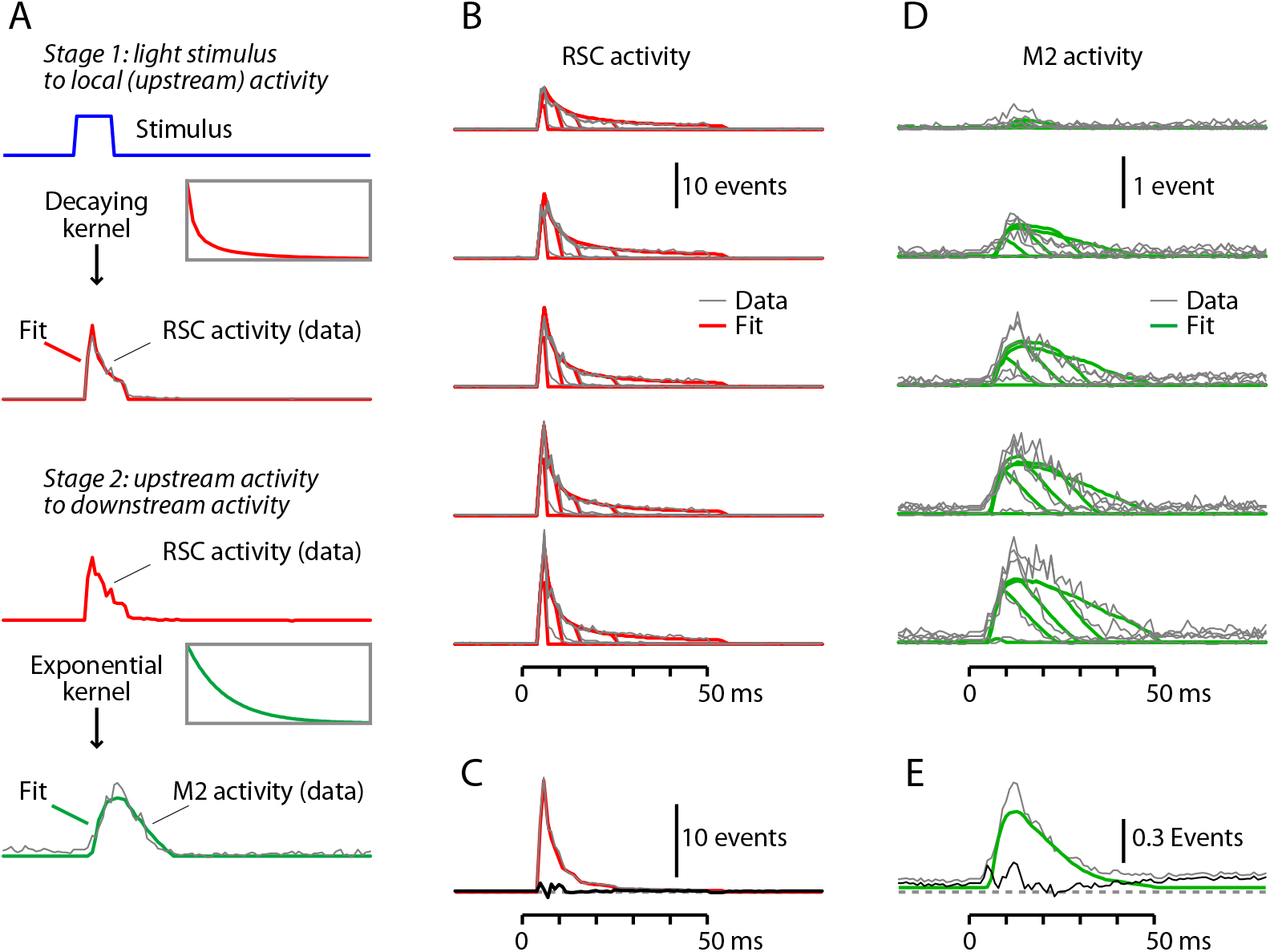
A simple two-stage model captures the major features of orthodromic driving. (A) Depiction of the modeling. The first stage is the conversion of light pulses into local activity in the RSC, which is modeled by convolving the step pulses of light with a step function scaled by a decay process. The second stage is the conversion of the upstream RSC activity into downstream M2 activity, which is modeled by convolving the RSC activity with an exponential process with a temporal lag. The models were fitted to the data over the 0-60 ms post-stimulus interval. See text for additional details. (B) The fitted RSC responses (red) were generated by modeling the light pulse→RSC transfer function as described in panel A. The AAV9 data traces (gray) are shown superimposed. (C) Plot of the residuals (black trace), calculated by subtracting the mean fitted traces (red) from the mean data traces (gray). (D) The fitted M2 responses (green) were generated by modeling the RSC→M2 transfer function as described in panel A. The data traces (gray) are shown superimposed. (E) Plot of the residuals (black trace), calculated by subtracting the mean fitted traces (green) from the mean data traces (gray).

Indeed, we found this first-stage model to produce good fits when we analyzed activity in the stimulated (RSC) area. We find that the model qualitatively describes the data, describing both its initial peak and its decay over time (**Fig. 5B,C**). The first-stage model does not capture the initial 3 ms (which were blanked), but since response amplitudes peaked later than this the model nevertheless captured the main features of the response. Moreover, the first-stage model has high *R*^2^ values on both the AAV9 (0.93) and the AAV1 (0.83) datasets. This suggests that the stimulation effect is largely described by a linear, essentially immediate translation of the ChR2-mediated depolarization into spiking activity, which decays rapidly.

Next, we developed a second-stage model for the corticocortical driving of M2 activity by RSC activity. Explorative data analysis revealed that activity in M2, the indirectly stimulated (downstream) area, could be fit well in terms of the activity of the stimulated area simply using thresholded activation, without an additional decay or adaptation process (**Fig. 5A**). We modeled this by convolving the upstream activity with an exponentially decaying kernel and applying a threshold. So this second-stage model has 4 parameters for gain, threshold, kernel time-constant, and baseline. These 4 parameters again seem intuitively necessary: the gain describes the strength of the corticocortically driven downstream activity; the threshold accounts for the inability of insufficiently strong upstream stimulation to produce any downstream activity; there is a slow transmission of information; and, there is non-zero baseline activity in the downstream area. Adding an explicit delay parameter to the second-stage model was not necessary: the combination of thresholding and slow stimulus integration sufficed to reproduce the experimentally observed the delay.

We found this second-stage model to produce good fits in the downstream (M2) area. We find that the model qualitatively describes the data, describing both its slow rise, and its subsequent decay over time (**Fig. 5D,E**). It also describes how in some conditions there is no activation whatsoever. This model also has high *R*^2^ values on both the AAV9 (0.70) and the AAV1 (0.65) datasets. The time constants of the fitted exponential kernels were on the order of a few tens of milliseconds (20 ms for AAV9 and 32 ms for AAV1 data), which may include contributions from many aspects, such as synaptic current and membrane time constants. It is also comparable to the time constants of fast spike adaptation in cortical excitatory neurons (La Camera et al., 2006; Wark et al., 2007; Suter et al., 2013). Thus the bulk of the response in the downstream area, M2, is described by linear integration of the input from the upstream area, RSC, with an effect that decays exponentially over time.

### Analysis of orthodromically driven responses

Next, we assessed whether the reduced amplitude of M2 responses (compared to upstream RSC activity, discussed above) was a consistent property across stimulus parameters. Plotting the response amplitudes in RSC and M2 for all 25 stimulus combinations (**Fig. 6A**) showed that these ranged widely but with a consistent relationship, substantially greater in RSC than in M2. The same pattern was observed for both viruses (factor of 4.7 for AAV9 and 6.8 for AAV1 experiments), even though absolute response amplitudes were generally stronger for AAV9 compared to AAV1 (1.5-fold for RSC responses and 2.1-fold for M2 responses; *p* < 10^−3^, sign test). Overall, the ‘driving ratio’, the ratio of the remotely driven activity in M2 relative to the locally driven activity in RSC, was ~0.2 (**Fig. 6B**). In other words, activity in the downstream area, M2, was generally about a fifth of that in RSC, across a wide range of stimulus parameters.

**Figure 6.**
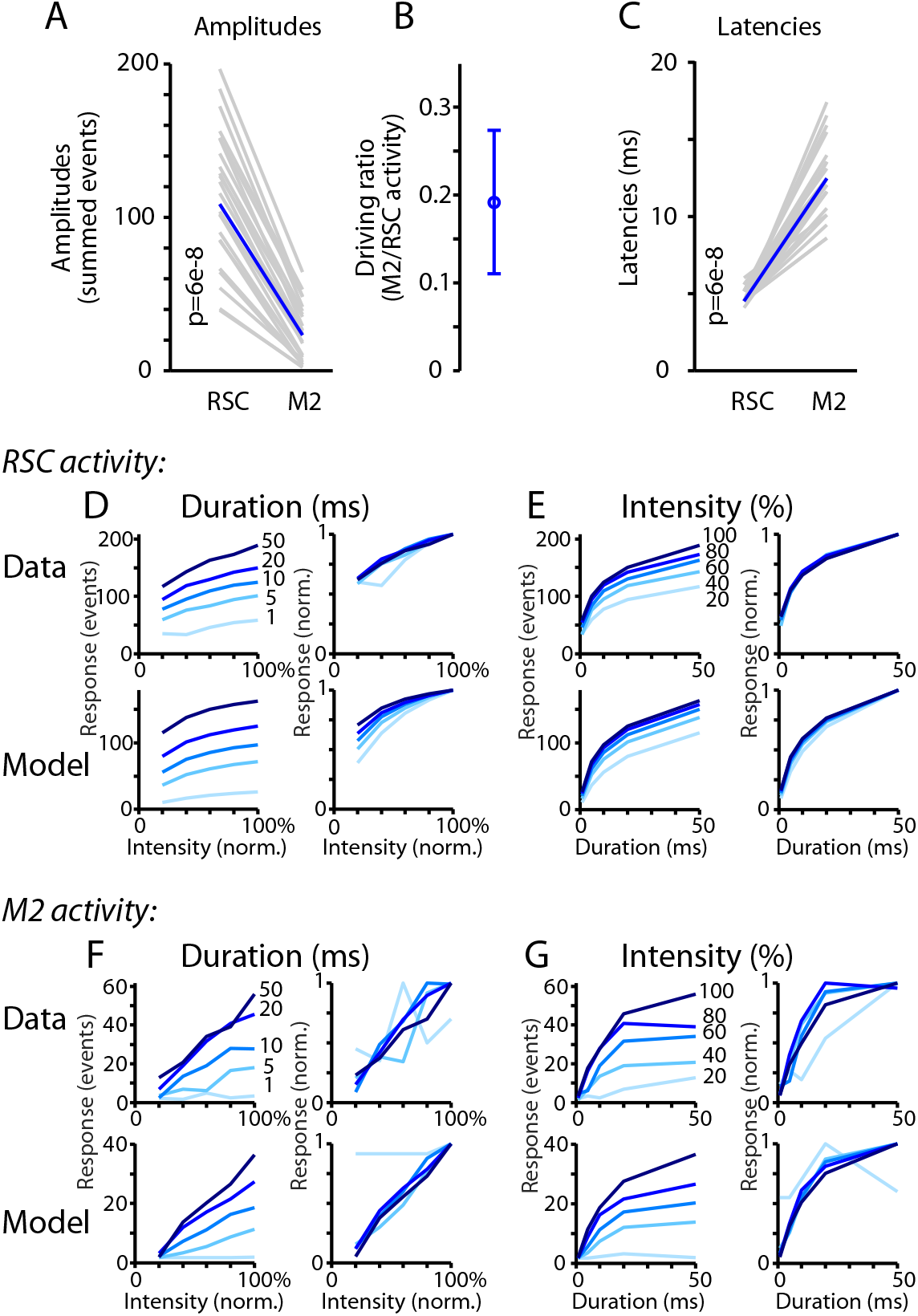
Analysis of orthodromically driven response amplitudes and latencies. (A) Amplitudes (calculated as the summed events) of the responses recorded on the RSC and M2 probes during RSC photostimulation, for each of the 25 combinations of stimulus intensity and duration (gray) along with the median values (blue), plotted for AAV9 experiments (see text for AAV1results). *P*-values calculated by 2-sided, paired sign test. (B) Driving ratios (defined as the ratio of activity generated locally in RSC over that generated remotely in M2) plotted as the median (across the 25 stimulus parameter combinations) ± m.a.d. (C) Same as panel A, but for latencies. (D) Dependence of RSC responses on stimulus intensity and duration. Left, top: For the RSC recordings, response amplitudes are plotted as a function of stimulus intensity; each line is for data recorded at constant stimulus duration, as indicated. Left, bottom: Same analysis, for the modeled responses. Right plots: same curves but peak-normalized. Response amplitudes grew approximately linearly with stimulus intensity. (E) Same analyses as panel D, but showing responses as a function of stimulus duration. Response amplitudes grew sub-linearly (approximately logarithmically) with stimulus duration. (F–G) Same as panels D-E, but for M2 recordings.

Of further importance to the interaction are latencies. These also showed a consistent relationship, with M2 responses peaking with a short delay after RSC responses (**Fig. 6C**). The same pattern was observed for both viruses (median latency of M2 response relative to RSC response of 8 ms for AAV9 and 7 ms for AAV1 experiments). In this case, unlike the absolute response amplitudes, the latencies of the responses in RSC and M2 did not differ significantly for AAV9 vs AAV1 (*p* > 0.05, sign test). In contrast to the amplitudes, the latencies were largely stimulus-independent.

Response amplitudes in both areas clearly varied systematically and substantially for different combinations of stimulus intensity and duration, but how? Plotting the RSC responses as a function of stimulus intensity showed a nearly linear dependence (**Fig. 6D**). In contrast, plotting the same RSC responses as a function of stimulus duration showed a sub-linear dependence (**Fig. 6E**). Applying the same analysis to the modeled traces gave qualitatively similar results (**Fig. 6D,E**, bottom row of plots). The M2 responses showed a similar, albeit noisier, set of patterns, with roughly linear intensity-dependence (**Fig. 6F**) and sub-linear duration-dependence (**Fig. 6G**). Applying the same analysis to the modeled traces again gave qualitatively similar results (**Fig. 6F,G**, bottom row of plots).

### Driving in reverse: antidromic propagation

The photoexcitability of ChR2-expressing axons (Petreanu et al., 2007) has previously been exploited in *in vivo* experiments to antidromically drive a trans-callosal corticocortical projection (Sato et al., 2014). Here, our experimental set-up (**Fig. 1**), by incorporating an optical fiber placed over the M2, allowed us to similarly drive the RSC→M2 projection in reverse, and thereby gain additional insight into signaling properties in this system. Characterization of antidromic optogenetic driving is additionally of technical interest both as an intended (e.g. (Sato et al., 2014)) or unintended and therefore potentially confounding effect of focal photostimulation in an area containing ChR2-expressing axons. Using the same labeling strategy (i.e., AAV-ChR2 in RSC) and recording (i.e., electrodes in both RSC and M2) arrangement, in the same experiments we also delivered photostimuli to M2 (via a second optical fiber) as a way to activate ChR2-expressing axons there (i.e., projecting from RSC) and thereby gain insight into the properties of antidromic signaling in the same RSC→M2 pathways (**Fig. 7A**).

**Figure 7.**
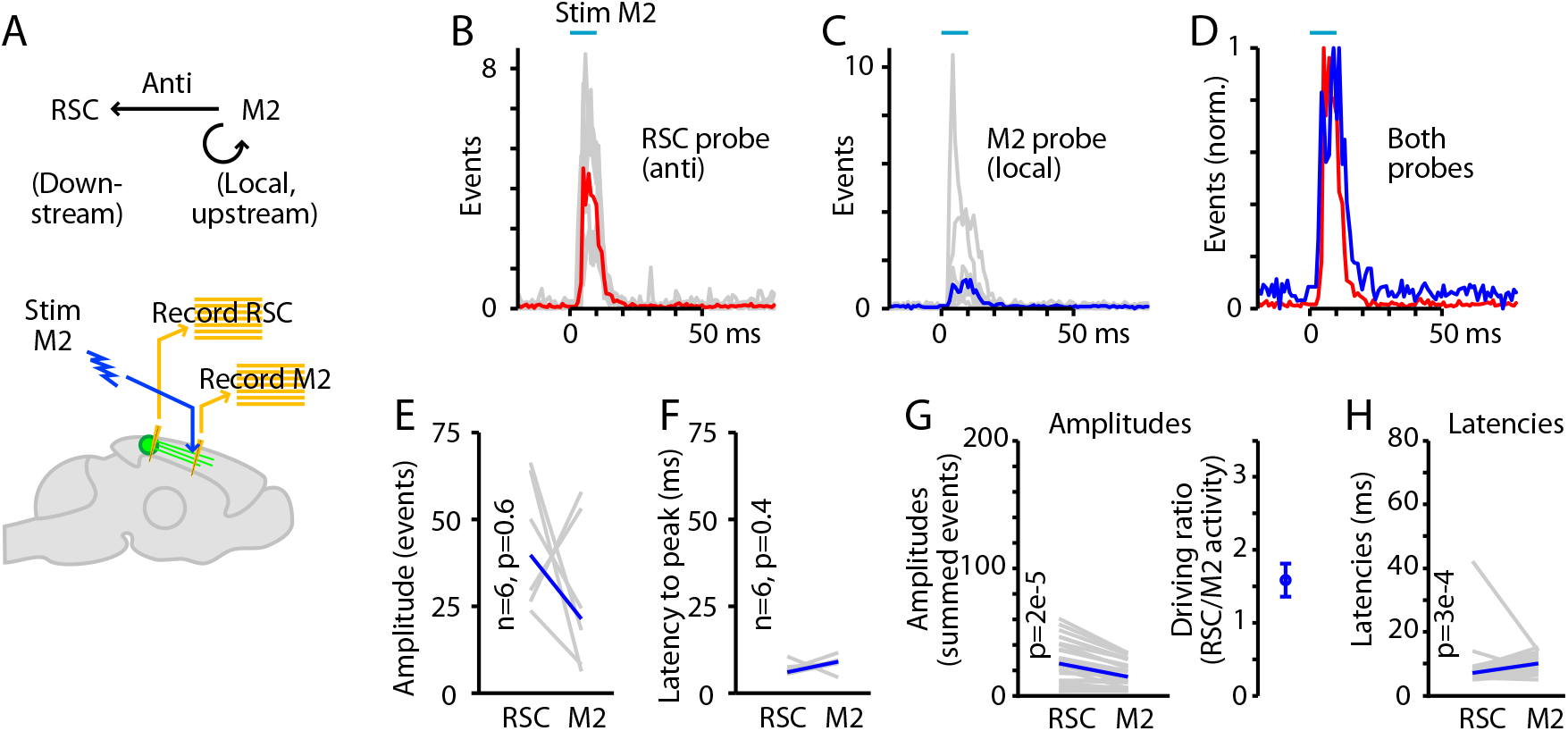
Driving in reverse: antidromic propagation. (A) Experimental paradigm: RSC neurons were infected with AAV to express ChR2, and photostimuli were applied to M2 (to stimulate axons of RSC neurons) while recording multi-unit activity in both M2 (locally driven) and RSC (antidromically driven). (B) Activity recorded on the RSC probe during RSC stimulation in an animal injected with AAV9-ChR2. Red trace is the median response across animals (traces for each animal shown in gray). (C) Activity recorded on the M2 probe during the same experiment. Blue trace is the median response across animals. (D) Overall activity on the RSC and M2 probes plotted together (peak-normalized). (E) Amplitudes of responses (summed events) recorded on the RSC and M2 probes, for the same stimulus parameter combination (10-ms duration, 100% intensity) used for the data shown in panels B-D, plotted for each experiment (gray) and as the median across animals (blue). *P*-value calculated by 2-sided, paired sign test. (F) Latencies (to peak) for responses recorded on the RSC and M2 probes (same stimulus). (G) Response amplitudes across all 25 stimulus parameter combinations (gray), with the overall median (blue), plotted for AAV9 experiments (see text for AAV1 results). Right: Driving ratio (defined as the ratio of activity generated locally in RSC over that generated remotely in M2) plotted as the median (across the 25 stimulus parameter combinations) ± m.a.d. Scaling of vertical axes is set to facilitate comparison to similar plots in Fig. 6. (H) Same, for latencies.

In particular, we wondered if antidromic activation would result in similar or different effects compared to orthodromic activation. Photostimulation in M2 (i.e., of the ChR2-expressing axons of RSC neurons) resulted in a short-latency, short-duration wave of antidromically generated activity in both RSC and a similar but smaller-amplitude wave of locally generated activity in M2 (**Fig. 7B-D**). Neither amplitudes nor latencies differed with antidromic activation for the standard (10-ms, 100% intensity) stimulus combination (**Fig. 7E,F**). However, across all stimulus combinations the response amplitudes were overall ~2-fold greater in RSC relative to M2 (**Fig. 7G**), contrasting with the reduced amplitude in the downstream area observed with orthodromic stimulation. Similar to orthodromic stimulation, absolute response amplitudes were generally stronger for AAV9 compared to AAV1 (2.6-fold for RSC responses and 3.8-fold for M2 responses; *p* < 10^−3^, sign test). Latencies in the two areas were indistinguishable with AAV1 and slightly delayed (by 3 ms) in M2 with AAV9 (**Fig. 7H**). Latencies in RSC were slightly shorter with AAV9 than AAV1 (by 2.5 ms; *p* < 10^−4^, sign test), but those in M2 were the same with the two viruses (*p* > 0.05, sign test). These results indicate that RSC axons forming this corticocortical projection can be robustly activated in M2, generating activity both locally in M2 and antidromically in RSC – which is in effect the ‘downstream’ area in this experimental configuration.

## DISCUSSION

We analyzed corticocortical signaling in the RSC→M2 pathway *in vivo* using optogenetic photostimulation and electrophysiology. Across a wide range of stimulus parameters, the downstream responses arrived rapidly and scaled systematically with the photo-evoked activity in the upstream area. We found that a simple model involving linear integration, delay, and thresholding could describe much of the data.

In using optogenetic photostimulation to analyze this circuit we did not attempt to mimic naturalistic activity patterns of the RSC but rather used this as a tool to drive the circuit in a highly precise, controlled manner (Miesenbock, 2009). This approach allowed us to selectively activate the upstream neurons in the RSC→M2 pathway, and to systematically vary the stimulus intensity and duration and assess whether and how response properties depended on input parameters. Focal optogenetic photostimulation differs fundamentally from non-specific methods for brain stimulation; extracellular electrical stimulation, for example, is inherently limited due to its nonspecificity, antidromic activation, and related issues (Histed et al., 2009; Joucla et al., 2012) and could not have been used to selectively study signaling in the RSC→M2 pathway.

Another artificial aspect of these experiments was the use of anesthesia, without which extensive parametric testing would have been challenging with head-fixed animals. Moreover, our studies focused on computational aspects of corticocortical population signaling, not how corticocortical signals relate to the high-dimensional aspects of behavior (Carandini, 2012). In awake animals, even “at rest” the patterns of functional connectivity in the cortex can be extremely complex and dissimilar to anatomical connectivity, whereas in anesthetized animals the structure-function correspondence is high (Barttfeld et al., 2015). Reduced complexity in the anesthetized state could reflect reduction of non-linearities of corticocortical signaling. Consistent with this possibility, ketamine anesthesia (used here) blocks NMDA receptors and other molecules involved in highly non-linear forms of signaling in cortical neurons (Antic et al., 2010; Sleigh et al., 2014). Our findings indicate highly linear signaling in the RSC→M2 corticocortical pathway in ketamine-anesthetized mice. Thus, one interpretation of our findings is that such signaling represents a kind of “ground state” for corticocortical communication in this pathway. We suggest that this simpler, linear mode of corticocortical signaling can serve as a robust basis for complex dynamic activity to emerge when non-linear mechanisms are active in the awake state. Further experiments will be needed to explore such speculations.

We found that a simple two-stage model captured the broad features of the data. At the upstream end, the conversion of light energy into local spiking activity in the upstream area (the RSC) could be described as a simple transfer function dominated by strong and rapid decay. The decay likely reflects primarily ChR2 desensitization, a property common to all ChR2 variants including the two used here (Nagel et al., 2003; Nagel et al., 2005; Lin et al., 2009). Additional components of the decay may have come from endogenous factors associated with the neurons and microcircuits in the locally stimulated area (e.g. GABA release from inhibitory interneurons, short-term synaptic depression). One potential application of this first-stage model of the local photoactivation process is that it could be used to design photostimuli that precisely compensate for the decay.

At the downstream end, the conversion of upstream activity (in RSC) into downstream activity (in M2) could be described by a simple exponential process with a brief delay, and no adaptation mechanism. Although a small non-linearity was included in the form of a threshold, the efficacy of the second-stage model suggests that corticocortical signaling is mostly linear. The efficacy of this model implies that corticocortical driving of downstream activity is highly scalable, and furthermore that adaptation (of corticocortical driving) is not a major factor in shaping the downstream response, at least on the short time scales (tens of milliseconds) studied here. However, some contribution of an adaptation process may be reflected in the early component of the responses, which tend to be larger than the fitted traces. Whether this simple model can describe corticocortical signaling in other inter-areal pathways remains to be determined, but similarities between our findings using optogenetic activation and related work in the visual system (e.g. (Carandini et al., 1997)) suggest this is plausible.

The scalability of corticocortical signaling observed here may be particular to the RSC→M2 pathway, or may represent a more general computational principle of cortical operation (Miller, 2016; Rolls, 2016). Although cortical circuit organization appears basically conserved, areas can also differ substantially in their quantitative properties (Harris and Shepherd, 2015). Corticocortical signaling in other pathways might therefore be expected to exhibit broadly similar scalability, but with pathway-specific differences in the details of spatiotemporal dynamics. The ability to capture both general and pathway-specific features of corticocortical signaling in a simple mathematical model suggests a utility of this approach both for theoretical approaches to cortical network modeling (Bassett and Sporns, 2017) and for neural engineering approaches in which closed-loop neural dynamics and behavioral control require predictive modeling (Grosenick et al., 2015). Further studies will be needed to test these speculations.

The downstream response latencies (~8 ms after upstream responses), together with the RSC-M2 inter-areal distance of ~2 mm and allowing for the timing of synaptic transmission (Sabatini and Regehr, 1999), implies a conduction speed for these RSC→M2 corticocortical axons on the order of 0.3 m/s, a typical value for thin unmyelinated cortical axons (Raastad and Shepherd, 2003). The consistency of the latency timing across different stimulus parameters suggests that the RSC→M2 circuit was activated in a similar manner independent of the particular activity level of the RSC neurons; in particular, this suggests that the M2 activity resulted from direct excitatory RSC input to M2 neurons, rather than polysynaptic pathways via posterior parietal cortex or anterior thalamus (Yamawaki et al., 2016) or hippocampus (Sugar et al., 2011). Had polysynaptic interactions been increasingly engaged by longer-duration stimuli, responses should have increased over time in both RSC and M2, not decreased as observed.

In addition to robust forward (orthodromic) activation, we found robust reverse (antidromic) corticocortical signaling in RSC→M2 circuits. Antidromic driving, evoked by stimulating in M2 the ChR2-labeled axons projecting from RSC, was notable for two distinct properties. First, photostimulation in M2 (of the ChR2-expressing axons of RSC neurons) generated even more activity in RSC than in M2, by a factor of ~2. Thus, the gain in this corticocortical circuit (ratio of downstream to upstream activity) appeared to be a property associated with the anatomical directionality of the projection (RSC→M2), rather than determined by the site of stimulation. The greater activity in RSC could reflect locally abundant axonal branches of the labeled RSC neurons. Second, the efficiency of information transmission appeared similar in either direction; i.e., a property associated with the site of stimulation rather than the anatomical directionality of the projection. Optogenetic antidromic activation has been previously exploited used as a way to selectively generate activity in an area (e.g. (Sato et al., 2014)). Our results thus not only provide an additional example of how a corticocortical pathway can be driven in reverse to remotely generate activity in an area of interest, but identify key similarities as well as differences compared to orthodromic driving.

Corticocortical signaling in the RSC→M2 pathway may be critical for conveying information from hippocampus-associated networks involved in spatial memory and navigation to cortical and subcortical networks involved in decision making and action planning and execution (Vann et al., 2009; Sugar et al., 2011; Yamawaki et al., 2016). Consistent with this, lesions of the RSC impair navigation without impairing either motor function or the ability to recognize navigational landmarks (Maguire, 2001), and RSC pathology can be an early and prominent feature of Alzheimer’s disease (Minoshima et al., 1997). Conversely, the RSC→M2 connectivity appears strengthened after damage to adjacent cortex in a mouse stroke model (Brown et al., 2009). Thus another potential application of experimental-theoretical paradigm developed here is to understand primary pathology and adaptive plasticity in corticocortical signaling in mouse models of disease.

## Acknowledgements

We thank C. Maguire and N. Bernstein for technical assistance, and D. Heeger and M. Landy for helpful discussions. Grant support: NIH (NINDS grant NS061963; NIBIB grant EB017695).

